# PRC1-dependent compaction of Hox gene clusters prevents transcriptional derepression during early *Drosophila* embryogenesis

**DOI:** 10.1101/250183

**Authors:** Thierry Cheutin, Giacomo Cavalli

**Affiliations:** Institute of Human Genetics, UMR 9002 of the CNRS and the University of Montpellier, 34396 Montpellier, France

**Author notes:** **Corresponding authors: Email:** (T.C.); (G.C.).

## Abstract

Polycomb-group (PcG) proteins are conserved chromatin factors that maintain the silencing of key developmental genes, notably the Hox gene clusters, outside of their expression domains [1-3]. Polycomb repressive complex 2 (PRC2) trimethylates lysine K27 of histone H3 [4], and PRC1 collaborates with PRC2 in gene silencing. Genome-wide studies have revealed large H3K27me3 chromatin domains bound by PcG proteins, and Polycomb domains fold into distinct nuclear structures [5-9]. Although PRC1 is involved in chromatin compaction [10-16], it is unknown whether PRC1-dependent transcriptional silencing is a consequence of its role on higher-order chromatin folding. This is because depletion of PRC1 proteins typically induces both chromatin unfolding and ectopic transcription, and ectopic transcription can open chromatin by itself. To disentangle these two components, we analysed the temporal effects of two PRC1 proteins, Polyhomeotic (Ph) and Polycomb (Pc), on Hox gene clusters during *Drosophila* embryogenesis. We show that the absence of Ph or Pc affects the higher-order chromatin folding of Hox clusters prior to ectopic Hox gene transcription, demonstrating that PRC1 primary function during early embryogenesis is to compact its target chromatin. During later embryogenesis, we observed further chromatin opening at Hox complexes in both Ph and Pc mutants, which was coupled to strong deregulation of Hox genes at this stage of development. Moreover, the differential effects of Ph and Pc on Hox cluster folding matches the differences in ectopic Hox gene expression observed in these two mutants, suggesting that the degree of Hox derepression in PcG mutants depends on the degree of structural constraints imposed by each PcG component. In summary, our data demonstrate that binding of PRC1 to large genomic domains during early embryogenesis induces the formation of compact chromatin to prevent ectopic gene expression at later time-points. Thus, epigenetic mechanisms such as Polycomb mediated silencing act by folding chromatin domains and impose an architectural layer to gene regulation.

## Main text

PcG proteins are conserved epigenetic components, essential for cell differentiation, which maintain gene silencing during development [3]. PcG proteins affect both Hox gene expression and chromatin compaction of *Drosophila* and mammalian Hox gene clusters [11, 12, 17]. In *Drosophila* embryos, the best characterized target genes of PcG proteins are Hox genes, which are grouped into two large chromatin clusters of 350 to 400 kb, covered by histone H3K27me3 [5, 18-20]. The Antennapedia complex (ANT-C) includes *lab*, *pb*, *Dfd*, *Scr* and *Antp*, which control the cell identity of anterior segments, whereas the bithorax complex (BX-C) contains *Ubx*, *abdA* and *AbdB* genes, which are responsible for the identity of posterior segments [21, 22]. PcG proteins form two main classes of complexes, PRC2, which is responsible for the deposition of H3K27me3 and PRC1. PRC1 complexes are further subdivided in canonical PRC1 (cPRC1), which contains the Polycomb protein that binds to H3K27me3 via its chromo domain [23], and non-canonical PRC1 complexes, which lack Polycomb and contain various other subunits [3]. In flies, Hox gene expression is regulated by the cPRC1, which is composed of Sce, Psc-Suz2 and two proteins that are specific components of this complex: Ph and Pc. The mechanism by which cPRC1 mediates gene silencing is not understood. One hypothesis suggests that these proteins compact chromatin to form facultative heterochromatin and prevent gene activation [24, 25]. However, since deletion of PRC1 proteins causes ectopic gene expression which can also open chromatin, a causal link is difficult to demonstrate. To distinguish whether higher-order chromatin folding precedes PRC1-dependent transcriptional silencing, we analysed the time-course of 3D chromatin compaction and Hox gene derepression in mutant embryos in which Ph or Pc have been deleted.

We first performed RNA fluorescence in situ hybridization (FISH) experiments in wild-type embryos to detect nascent transcripts of eight *Drosophila* Hox genes, using probes recognizing the first introns of those genes. As expected, our results show sequential expression of *lab*, *pb*, *Dfd*, *Scr, Antp*, *Ubx*, *abdA* and *AbdB* along the anteroposterior axis [21, 26] at the germ band elongated stage (3:50-7:20 after fertilization) (Extended Data Fig. 1a-f), which is conserved throughout *Drosophila* embryogenesis (Extended Data Fig. 2). We then performed a series of immuno-DNA FISH experiments in embryos at the germ band elongated stage to address 3D chromatin compaction of Hox clusters. We measured 3D distances between FISH spots for *Ubx*, *abdA* and *AbdB* of the BX-C cluster or *lab*, *Scr* and *Antp* of the ANT-C cluster. The variation of these inter-spot distances along the anteroposterior axis shows that transcription of each Hox gene correlates with the opening of its corresponding chromatin region, whereas the silenced portion of Hox complexes remains condensed (Extended Data Fig. 1g-j).

We then performed RNA FISH experiments in mutant embryos that were deficient in the Ph (*ph^del^*) and Pc (*Pc^XT109^*) subunits. In both mutants, Hox gene derepression started in a few cells, and the proportion of cells with derepression increased during later embryogenesis (Fig. 1f-k; Extended Data Fig. 2-3). *Ubx* was the first gene of the BX-C cluster to be expressed ectopically in the anterior PS of *ph^del^* embryos, whereas derepression of *abdA* and *AbdB* started later (Fig. 1a, f-h). Similarly, *Antp* was the first ANT-C gene to become derepressed in the head of both mutant embryos (Fig. i-k), whereas the others were derepressed at later stages (Extended Data Fig. 2-3). In both mutants, ectopic expression of each Hox gene depended on its position along the anteroposterior axis. For example, the number of cells showing ectopic *Ubx* expression decreased from PS4 to the anterior PS (Fig. 1a-b; Extended Data Fig. 2q and r), and *Scr* derepression was enhanced in the posterior PS compared with that in the head (Fig. 1j). Taken together these results indicated that Ph and Pc do not elicit a general silencing function of all Hox genes. Indeed, although both proteins bind every Hox gene where they are repressed [27], loss of Hox gene silencing caused by Ph or Pc deletion was heterogeneous for different Hox genes. In addition, ectopic Hox gene transcription was generally stronger and started earlier in *ph^del^* embryos than in *Pc^XT109^* embryos (Extended Data Fig. 2). For example, *Ubx* and *Antp* derepression occurred as early as 3:50-4:50 after fertilization in *ph^del^* embryos, but only after the 4:50-6:00 stage of embryonic development in Pc^XT109^ (Fig.1 f, k). In contrast, *AbdB* was derepressed earlier in *Pc^XT109^* mutants than in *ph^del^*, especially in PS7-PS12 (Fig. 1h; Extended Data Fig. 2w-x). This suggests that different cPRC1 subunits play specific roles on their target chromatin.

**Figure 1:**
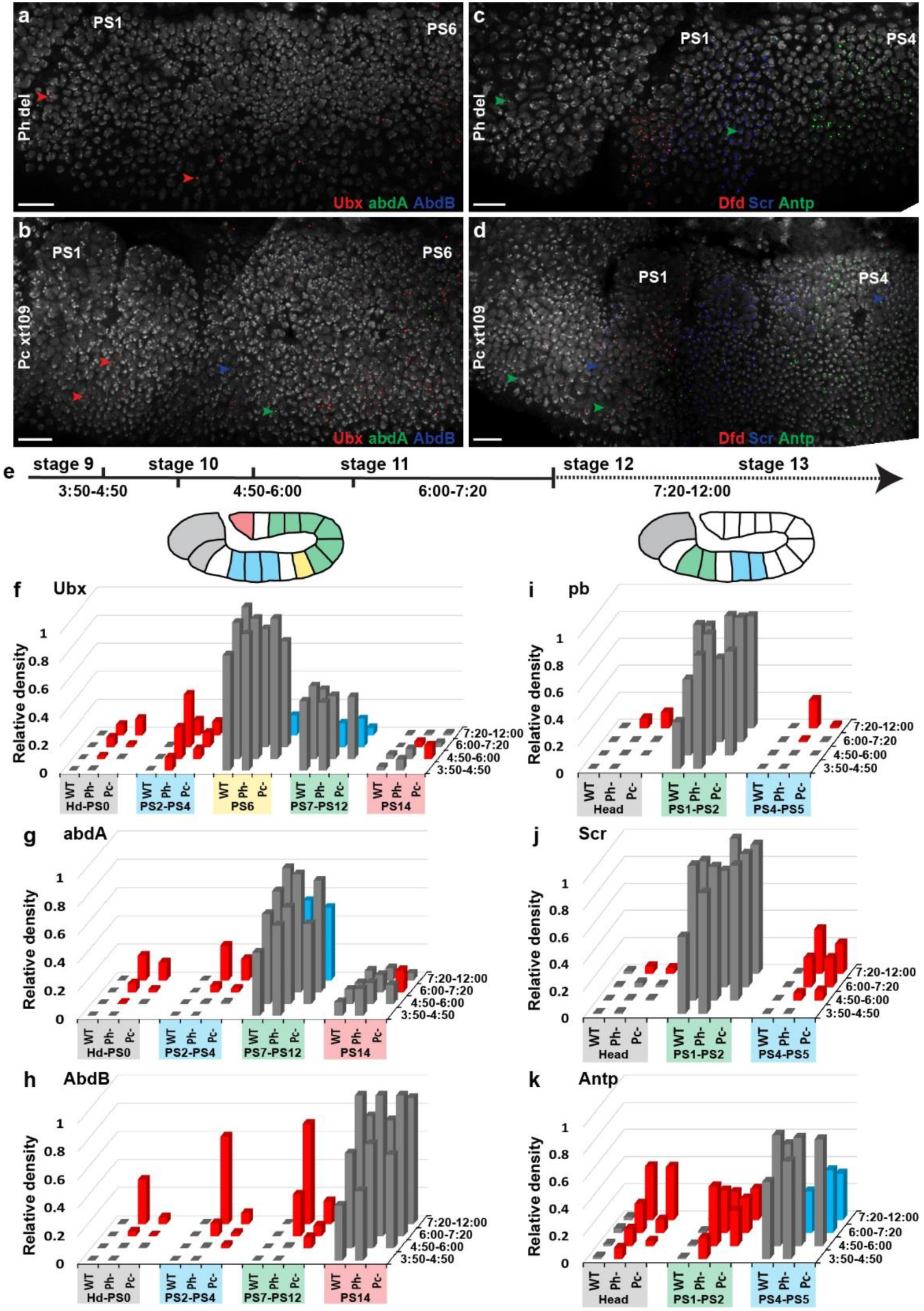
Timing of Hox gene derepression in *Ph^del^* and *Pc^XT109^* embryos. **a-d,** RNA FISH images illustrating the earliest ectopic Hox gene expression observed in *ph^del^* (a-b) and *Pc^XT109^* (c-d) embryos. Arrowheads indicate few cells showing Hox gene transcription outside of their domains of expression (*Ubx*: red in a, b; *abdA*: green in b; *AbdB*: blue in b; *Antp*: green in c, d and Scr: blue in d). Scale bars, 20 μm. **e,** Embryos were grouped into four classes based on their developmental stage, which depended on the duration of their development after fertilization at 25°C. **f-k,** Relative densities of RNA FISH spots corresponding to *Ubx* (f), *abdA* (g), *AbdB* (h), *pb* (i), *Scr* (j) and *Antp* (k) expression measured in WT, *Ph^del^* and *Pc^XT109^* embryos during development. For simplicity, PSs wherein Hox genes of BX-C (f-h) and ANT-C (i-k) behave similarly were grouped (Complete Data are shown in Extended Data Figure 2). Red columns indicate where and when Hox gene transcription was significantly (Mann-Whitney U test, *P* < 0.01) ectopically expressed in the mutants compared to WT embryos, whereas the blue columns show significant (Mann-Whitney U test, *P* < 0.01) downregulation of Hox gene transcription.

We then tested the possibility that cPRC1 mediates direct compaction of Hox clusters to prevent ectopic transcription. We reasoned that, if it does so, mutations in cPRC1 components would affect Hox compaction prior to any detectable transcriptional activation. To test this hypothesis, we performed DNA FISH experiments to monitor BX-C and ANT-C chromatin folding in the cell nuclei of *ph^del^* and *Pc^XT109^* embryos. At the 3:50-4:50 stage after fertilization, distances between *Ubx*-*abdA*, *abdA*-*AbdB* and *Ubx*-*abdB* were significantly increased in the head and PS0 of *ph^del^* mutant embryos compared to those in control embryos (Fig. 2a-c), whereas neither *Ubx*, *abdA* nor *AbdB* were derepressed (Fig. 1f-h). Similar effects were observed in PS2-PS4 (Fig. 2g-i) where the derepression of *Ubx* occurred in only a few cells of *ph^del^* embryos (Fig. 1f). Despite a weaker effect of Pc on BX-C folding, *Pc^XT109^* mutant embryos displayed significantly greater *Ubx*-*abdA*, *abdA*-*AbdB* and *Ubx*-*abdB* distances than those of control embryos in PS2-PS4 at the 3:50-4:50 stage of development (Fig. 2g-i), without derepression of *Ubx*, *adbA* and *AbdB* (Fig. 1f-h). Similarly, Ph and Pc were both required to globally compact ANT-C in the heads of embryos from the 3:50-4:50 stage after fertilization (Fig. 3a-c), whereas *Antp* was the only Hox gene of the ANT-C cluster to be derepressed, and this occurred only in *ph^del^* mutants (Fig. 1i-k). These results show that in PSs where every Hox gene of one complex is repressed, the first effect of Ph and Pc on Hox clusters folding can be detected before ectopic Hox gene transcription and affected whole Hox complexes, whereas the first effects on Hox genes derepression affected a minority of the Hox genes.

**Figure 2:**
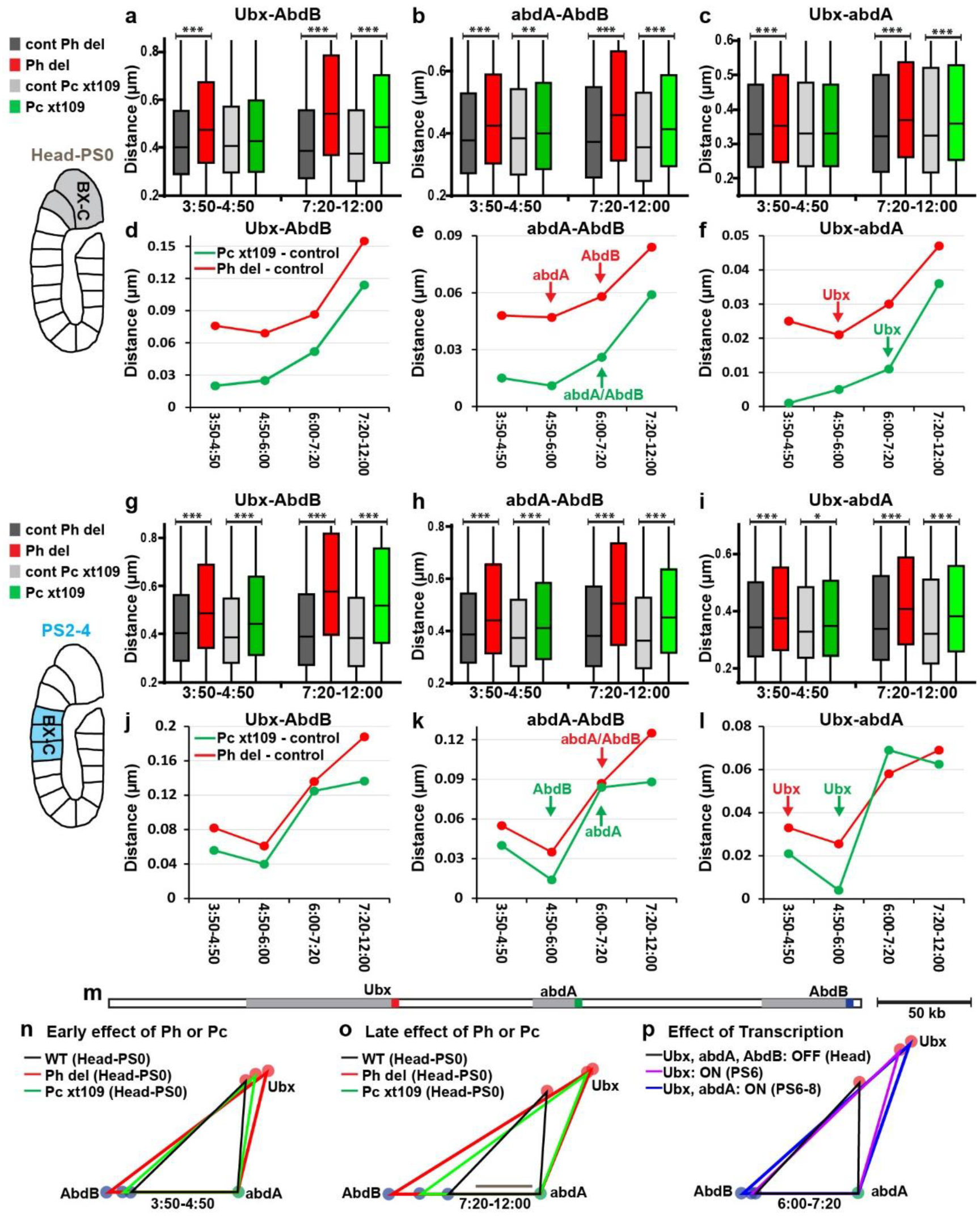
Ph and Pc are required to compact repressed BX-C before ectopic Hox gene transcription. **a-c; g-i;** Box plots displaying distributions of the distances *Ubx*-*AbdB* (a; g), *abdA*-*AbdB* (b; h), *Ubx*-*abdA* (c; i), in the head-PS0 (a-c), and in PS2-PS4 (g-i) during early and late embryogenesis. Distances were measured in the cell nuclei of *ph^del^* embryos (red) and their respective controls (dark grey) or *Pc^XT109^* embryos (green) and their respective controls (light grey). The lower and upper bounds of the coloured rectangles correspond to the first and third quartiles, whereas the middle bars show the median distances. The black lines indicate significant differences between the mutants and their respective control embryos (Mann-Whitney U test, * *P* < 0.05; ** *P* < 0.01; *** *P* < 0.001). **d-f; j-l;** Difference between the median distances *Ubx*-*AbdB* (d; j), *abdA*-*AbdB* (e; k), *Ubx*-*abdA* (f-l) measured in *ph^del^* and their control embryos (red) or in *Pc^XT109^* and their control embryos (green), during embryonic development in head-PS0 (d-f) and PS2-PS4 (j-l). Arrows indicate when a Hox gene is firstly ectopically expressed in *ph^del^* (red) or *Pc^XT109^* (green) embryos. **m,** Schematic linear representation of BC-X showing the DNA FISH probe positions. **n-p**, Comparison of the effects of Ph and Pc on the folding of BX-C during early (n) and late embryogenesis (o) with the openings induced by Hox gene transcription (p). Scale bars, 100 nm.

To compare Ph and Pc, we plotted the effect of *ph* or *Pc* deletions on distances measured within the BX-C (Fig. 2d-f; j-l) or the ANT-C (Fig. 3d-f). After the 4:50-6:00 stage, the effect of both proteins on BX-C folding progressively increased (Fig. 2d-f; j-l) with a timing matching the ectopic expression of *abdA* and *AbdB* (Fig. 1g-h). 7:20-12:00 hours after fertilization, both mutant embryos showed a stronger opening of the BX-C, in the head-PS0 (Fig. 2a-c) and PS2-PS4 (Fig. 2g-i).. To summarize these effects, we plotted the three median distances between the promoters of *Ubx*, *abdA* and *AbdB* (Fig. 2m-p) or between *lab*, *Scr* and *Antp* (Fig. 3g-j). During early embryogenesis, the effects of the loss of Ph and Pc on Hox cluster folding were significant (Fig. 2a-c, n; 3a-c, h and Extended Data Fig. 4a-f), although a stronger decompaction of Hox clusters was observed in later embryogenesis (Fig. 2a-c, o; 3a-c, i and Extended Data Fig. 4g-l), consistent with strong ectopic Hox expression. These late effects coincide with the pattern of distance changes observed during physiological Hox activation in the appropriate PSs (Fig. 2p, 3j). Therefore, the strong effects on Hox distances observed in late development in the mutants is most likely due to the effect of ectopic transcription. Taken together, these results demonstrate that the loss of PRC1 prevents the condensation of Hox clusters prior to any transcriptional derepression. Thus, chromatin opening in the mutants is not a consequence of transcription, suggesting that the primary function of PRC1 is to establish a compact architecture in cells where Hox loci are silenced.

**Figure 3:**
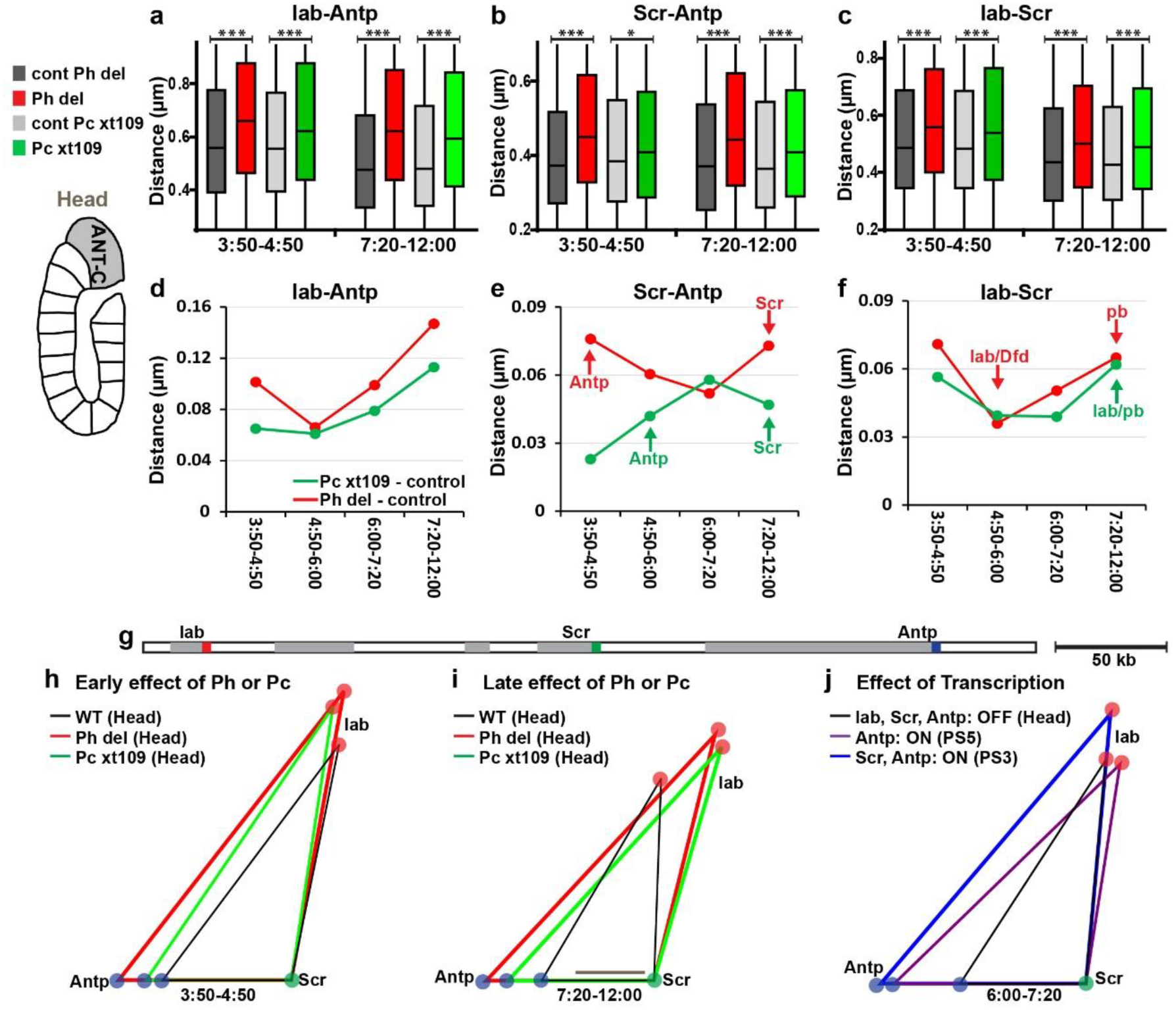
Ph and Pc are required to compact repressed ANT-C before ectopic Hox gene transcription. **a-c,** Box plots displaying distributions of the distances: *lab*-*Antp* (a), *Scr*-*Antp* (b) and *lab*-*Scr* (c) in the head during early and late embryogenesis. Distances were measured in the cell nuclei of Ph^del^ embryos (red) and their respective controls (dark grey) or Pc^XT109^ embryos (green) and their respective controls (light grey). The lower and upper bounds of the coloured rectangles correspond to the first and third quartiles, whereas the middle bars show the median distances. The black lines indicate significant differences between the mutants and their respective control embryos (Mann-Whitney U test, * *P* < 0.05; ** *P* < 0.01; *** *P* < 0.001). **d-f,** Difference between the median distances *lab*-*Antp* (d), *Scr*-*Antp* (e) and *lab*-*Scr* (f) measured in *ph^del^* and their control embryos (red) or in *Pc^XT109^* and their control embryos (green) during embryonic development in head. Arrows indicate when a Hox gene is firstly ectopically expressed in *ph^del^* (red) or *Pc^XT109^* (green) embryos. **g,** Schematic linear representation of ANT-C showing the DNA FISH probe positions. **h-j,** Comparison of the effects of Ph and Pc on the folding of ANT-C inside cell nuclei during early (h) and late embryogenesis (i) with the openings induced by Hox gene transcription (j). Scale bars, 100 nm.

Finally, we investigated the consequences of *Pc-* or *ph*-null mutations on Hox loci in regions in which they are actively transcribed in WT embryos. No effect on Hox genes transcription in these regions were revealed by RNA FISH analysis during early embryogenesis (3:50-6:00 after fertilization) (Fig. 1f-k). In PS9-PS12, where *Ubx* and *abdA* are expressed but *AbdB* is not, no significant effect on the *Ubx*-*abdA* distance was observed. However, as expected, the *abdA*-*AbdB* distance was increased in both *ph^del^* and *Pc^XT109^* embryos compared to control embryos (Fig. 4a-b). Conversely, the distance *abdA*-*AbdB* was not increased in PS14 where *AbdB* is expressed, while the distance *Ubx*-*abdA* increased in both mutants during embryogenesis (Fig. 4c-d). Similarly, the lack of Ph or Pc did not increase the distance *Scr*-*Antp* in PS4-PS5, where *Antp* is expressed, while the distance *lab*-*Scr* was significantly increased in the same region (Fig. 4e-f). These results demonstrate that Pc and Ph compact chromatin fibres encompassing Hox genes only in cells in which they are repressed (Extended Data Fig. 5-6).

**Figure 4:**
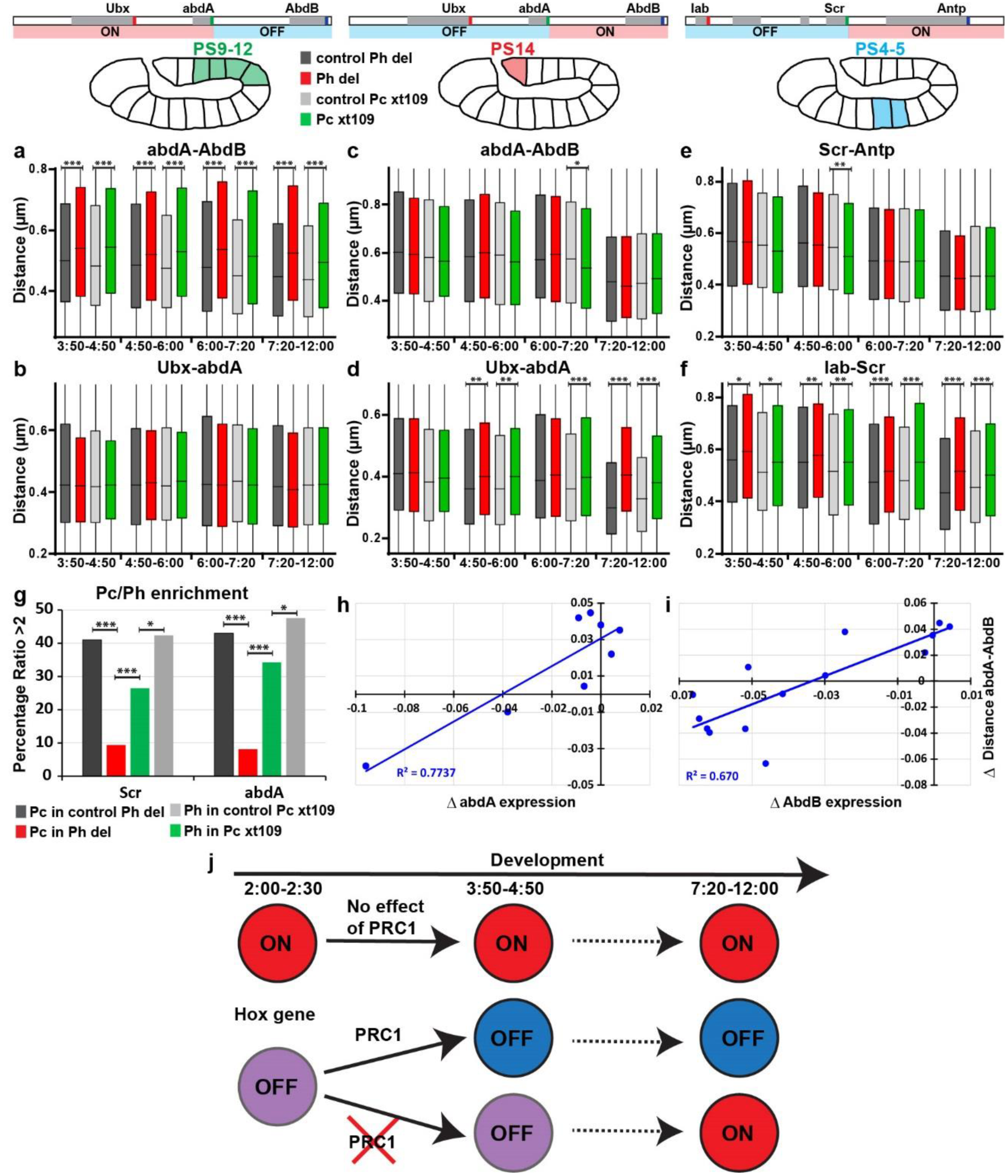
cPRC1 only compacts chromatin in the repressed regions of the BX-C and ANT-C domains. **a-f,** Box plots displaying distributions of the distances *abdA*-*AbdB* (a; c), *Ubx*-*abdA* (b; d), *Scr*-*Antp* (e) and *lab*-*Scr* (f) measured in the cell nuclei of *Ph^del^* embryos (red) and their respective controls (dark grey) or *Pc^XT109^* embryos (green) and their respective controls (light grey). Measurements were made in PS9-12 (a-b), PS14 (c-d), and PS4-5 (e-f) during development. The lower and upper bounds of the coloured rectangles correspond to the first and third quartiles, whereas the middle bars indicate the median distances. **g,** Charts presenting the percentage of DNA FISH spots with Pc or Ph enrichment more than two times the average intensity inside the cell nuclei. Significant differences are indicated (Mann-Whitney, U test, * *P* < 0.05; ** *P* < 0.01; *** *P* < 0.001). **h-i,** Scatterplots showing correlations between the differential effect of Ph and Pc on the distance *abdA*-*AbdB* at 3:50-4:50 and the differential effect of Ph and Pc on *AbdB* or *abdA* expression at 4:50-6:00. Each point corresponds to one PS where *abdA* (h) or *AbdB* (i) are repressed in WT embryos. **j,** Schematic diagram summarizing the effects of PRC1 on Hox gene folding and transcription. Circles represent silenced (OFF) or transcribed (ON) chromatin associated with Hox genes (red, open; purple, partly compact; blue fully compact). cPRC1 has no effect when Hox genes are expressed and chromatin is open. When Hox genes are repressed, cPRC1 compacts their chromatin during early embryogenesis and they will remain silenced. Without cPRC1, this compaction cannot occur and silenced Hox genes might become subsequently transcribed.

Interestingly, Pc was found to have a weaker effect than Ph on Hox clusters folding in the PS where every Hox gene of each complex is repressed (Fig. 2-3). Consistent with a difference between these two cPRC1 subunits, the nuclear Pc distribution became diffuse in *ph^del^* embryos, whereas Ph still accumulated in nuclear foci in *Pc^XT109^* embryos (Extended Data Fig. 7a-c). Immuno-FISH experiments using anti-Ph and anti-Pc antibodies and FISH probes recognizing either *abdA* or *Scr* showed that Ph protein still accumulated at *abdA* and *Scr* loci in Pc^XT109^ embryos. This is in contrast to the nuclear distribution of Pc, which did not accumulate on the same genes in *ph^del^* embryos (Fig. 4g; Extended Data Fig. 7d-e). In contrast, the effect of Pc and Ph on the distance *abdA*-*AbdB* is similar in posterior PSs (Fig. 4a; Extended Data Fig. 5b, e). For each PS where *abdA* or *AbdB* is repressed in WT embryos, we calculated the difference of *abdA* or *AbdB* expression between *ph^del^* and *Pc^XT109^* mutant embryos at stage 4:50-6:00. Scatterplots between these latter values and the difference of distance *abdA*-*AbdB* between *ph^del^* and *Pc^XT109^* mutant embryos at stage 3:50-4:50 showed correlations between chromatin opening and later ectopic transcription (Fig. 4h-i), suggesting a causal link between chromatin condensation and gene silencing.

Taken together, these results demonstrate that cPRC1 compacts Hox clusters via the formation of higher-order chromosome structures during early *Drosophila* embryogenesis (Fig. 4j). The general effect on chromatin compaction contrasts with the transcriptional effects observed in the absence of these proteins, with ectopic transcription occurring later than chromatin opening, with timing and localizations that are specific to each Hox gene. It can be speculated that these differences reflect the relative abundance and target site affinity for cognate transcription factors. Moreover, the roles of the Ph and Pc proteins in the formation of PRC1 foci and Hox gene silencing are not equivalent, with Ph showing strongest effects, except on the *abdA* and *AbdB* region of the BX-C. Although the molecular reasons for this difference are not known, one possible explanation is that, in the absence of Pc and of its chromo domain, cPRC1 might only lose its anchoring to H3K27me3 while retaining some of its ability to bind target regulatory elements and mediate their clustering through oligomerization [8, 17, 28] (Extended Data Fig. 7f-g) [29]. The comparatively strong effect of *Pc* deletion on the *abdA-AbdB* region of the BX-C locus might be explained by the fact that H3K27me3 levels are highest in this region compared to all others [27]. On the other hand, in the absence of Ph, cPRC1 might either dissolve or it might lose its ability to bind its target loci, therefore inducing strong decompaction throughout the BX-C and ANT-C and, consequently, a generally stronger loss of silencing than *Pc* mutants. Further studies will be required to elucidate this point and the mechanism of PRC1-mediated silencing at other genes. In particular, it will be interesting to test whether the role of PRC1 in chromatin condensation is only predominant at large Polycomb domains containing many PRC1 binding sites and whether the mechanisms of silencing differ at smaller target loci, both in *Drosophila* and in mammals.

## Acknowledgements

We thank Montpellier Resources Imaging facility MRI-IGH for microscopy support. We thank Satish Sati and Frédéric Bantignies for critical reading of the manuscript. G.C. Research in the laboratory of G.C. was supported by grants from the European Research Council (ERC-2008-AdG No 232947), the CNRS, the FP7 European Network of Excellence EpiGeneSys, the European Union’s Horizon 2020 research and innovation programme under grant agreement No 676556 (MuG), the Agence Nationale de la Recherche, the Fondation pour la Recherche Médicale, INSERM, the French National Cancer Institute (INCa) and the Laboratory of Excellence EpiGenMed.

## Author contributions

T.C., and G.C. initiated and led the project. T.C performed experiments and data analysis. T.C. and G.C. interpreted and discussed the data and wrote the manuscript.

